# A tumor suppressor role of the miR-15b/16-2 cluster in T-cell acute lymphoblastic leukemia

**DOI:** 10.1101/2025.07.07.663472

**Authors:** Sara González-García, María J. García-León, Marina García-Peydró, Patricia Fuentes, Juan Alcain, Enrique Martín-Gayo, Carlo M. Croce, Ramiro Garzón, María L. Toribio

## Abstract

T-cell acute lymphoblastic leukemia (T-ALL) is an aggressive hematological malignancy arising from the neoplastic transformation of immature T cells during their development in the thymus. Deciphering the developmental programs whose dysregulation leads to T-ALL pathogenesis is critical for the development of novel targeted therapies, which remain an urgent unmet need for the treatment of this disease. MicroRNAs (miRNAs) have emerged as key post-transcriptional regulators of numerous physiological processes and cancer. However, the specific role of miRNAs in human T-cell development and T-ALL pathogenesis remains largely unexplored. In this study, we comprehensively evaluated miRNA expression profiles across human T-cell development by microarray analysis and identified a dynamic expression pattern of miR-16-2, which is upregulated across early pre-T cell proliferative stages up to the resting stage of immature thymocytes immediately preceding TCRαβ expression, and decreased thereafter. We confirmed the coordinated regulation of miR-15b expression, consistent with the reported clustered genomic location of both miRNAs. Notably, functional studies identified the miR-15b/16-2 cluster as a negative regulator of early thymocyte proliferation, and showed that overexpression of miR-15b/16-2 in T-ALL cells impaired leukemic growth *in vitro* and tumor progression in patient-derived xenotransplantation assays. Mechanistically, miR-15b/16-2 expression represses the genes encoding BCL-2 and CYCLIN D3, leading to T-ALL apoptosis and cell cycle dysregulation, with an accumulation of G0-phase cells and a defective transition to the G2/M phase. Overall, these findings support a novel function for miR-15b/16-2 as tumor suppressors in T-ALL, highlighting their role as promising targets for T-ALL therapy.

**KEY POINTS:**

- miR-15b/16-2 overexpression impairs the proliferation of human thymic progenitors leading to defective T-cell production.
- miR-15b/16-2 represses BCL-2 and CYCLIN D3 expression and impairs tumor progression in human T-ALL, revealing its role as tumor suppressor.

## INTRODUCTION

Dysregulation of T-cell maturation within the thymus can lead to the malignant transformation and developmental arrest of immature T cells, ultimately giving rise to aggressive T-cell acute lymphoblastic leukemia (T-ALL).^1^ While current therapies have significantly improved patient outcomes,^2,3^ 20-50% of patients, most notably adults, still experience relapse or develop refractory disease (r/r T-ALL), which is associated with a poor prognosis.^3–5^ Therefore, there is an urgent need of novel targeted therapies for these patients. Achieving this goal requires a deeper understanding of the molecular pathways that link aberrant T-cell development to leukemogenesis. Human T-cell development proceeds from lymphomyeloid CD34^hi^ CD44^hi^ CD1a^-^ early thymic progenitors (ETPs), which commit into pre-T cells (CD34^+^ CD44^lo^ CD1a^+^) capable of generating both αβ and ψ8 T cells. Development along the αβ lineage involves transition through CD4^+^ CD3^-^ immature single positive (CD4ISP) and CD4^+^ CD8^+^ double positive (DP) thymocytes. At this stage, successful TCRβ rearrangement leads to expression of the pre-TCR (CD3^lo^ pre-TCR^+^ TCRαβ^-^), which initiates β-selection, a critical process that triggers cellular expansion, pre-TCR downregulation, and progression into resting DP CD3^-^, leading to DP TCRαβ^+^ thymocytes. Subsequent positive and negative selection result in the generation of mature CD4^+^ or CD8^+^ single positive (SP) T cells expressing the TCRαβ.^6^ Identifying the molecular mechanisms that govern cell proliferation and survival throughout this complex developmental sequence is essential for uncovering novel T-ALL targeted therapies.

MicroRNAs (miRNAs) are small non-coding RNAs that repress gene expression by targeting messenger RNAs (mRNAs) for degradation or inhibiting translation.^7^ They have emerged as key regulators of multiple physiological processes and pathologies, including cancer.^8,9^ The importance of miRNAs in T-cell development was initially evidenced in mice, where conditional deletion of *Dicer*, an RNase III-like enzyme essential for miRNA biogenesis, resulted in impaired T-cell generation.^10,11^ Additional reports have described the contribution of individual miRNAs to distinct murine T-cell developmental stages, including β-selection-induced proliferation, and positive and negative selection.^12-15^

In humans, miRNA expression analyses across T-cell development are limited to DP and SP stages,^12,13^ but the miRNA signatures associated to earlier stages are largely unknown. Moreover, studies addressing the impact of dysregulated miRNA expression in developing human T cells are still missing, although aberrant miRNA expression during T-cell development is thought to contribute to T-ALL pathogenesis.^14–16^ Indeed, miRNAs can act either as oncogenes, ^17–19^ or as tumor suppressors in T-ALL.^20,21^ However, the mechanisms involved in miRNA-mediated development of this malignancy remain poorly understood.

Here, we conducted a comprehensive analysis of miRNA expression during human intrathymic T-cell development, identifying a coordinated expression of miR-15b and miR-16-2, members of the miR-15b/16-2 cluster. The observed dynamic expression pattern pointed to a tight developmental control of miR-15b/16-2 levels during early thymocyte proliferation, suggesting their potential impact in T-ALL pathogenesis. Accordingly, miR-15b/16-2 overexpression impaired T-ALL cell proliferation and progression, both *in vitro* and *in vivo*. These findings identify miR-15b/16-2 as novel tumor suppressors in T-ALL, highlighting their potential relevance as therapeutic targets for this malignancy.

## METHODS

### Isolation of human intrathymic T-cell populations and primary T-ALLs

Experiments were performed and human samples were obtained in accordance with approved guidelines established by Local Bioethics Committees (Spanish Research Council; Hospital Universitario 12 de Octubre and Hospital Universitario Niño Jesús, Madrid, Spain), and with the Declaration of Helsinki. Human thymocyte populations at sequential T-cell maturation stages (**Supplemental Table 1**) were isolated from pediatric thymus samples (1 month to 4 years) as previously described.^22^ Patient-derived T-ALL lymphoblasts were isolated from peripheral blood (PB) or bone marrow (BM) samples obtained at diagnosis as described.^23^

### Cell lines and *in vitro* cultures

Human T-ALL cell lines JURKAT, HPBALL, SUPT1 (American Type Culture Collection, ATCC), and CUTLL1 (kindly provided by Prof. Ferrando, Columbia University Irving Medical Center) were cultured in RPMI 1640 (Lonza) plus 10% fetal bovine serum (FBS, Gibco). Patient-derived T-ALL cells were cultured onto OP9 stroma (ATCC) in α-MEM (Gibco) plus 20% FBS, and recombinant human (rh) IL-7 (200 IU/mL, National Institute of Biological Standards and Controls -NIBSC-). HEK293T (ATCC) cells were cultured in DMEM plus 10% FBS.

### miRNAs microarray analysis

Total RNA was isolated from intrathymic T-cell populations (**Supplemental Table 1**) using TRIzol (Invitrogen). RNA integrity was confirmed using a 2100 Bioanalyzer (Agilent Technologies). Five μg of RNA was hybridized onto a custom miRNA microarray chip (OSU-CCC, Version 4.0), which includes 474 human miRNA genes spotted in duplicates.^24–26^ Microarray images were analyzed using GenePix Pro v.6.0 (Molecular Devices). Datasets were generated from three biological replicates per cell type, obtained from different thymus samples. Average values were background subtracted; log2-transformed, and normalized using quantiles normalization (Bioconductor). miRNAs were retained if present in at least 20% of samples. Data analyses were performed using BRB-ArrayTools v3.6.0, and R v2.3.1 (R Foundation for Statistical Computing) with a *P*-value <0.05. Hierarchical clustering was performed using average-linkage based on Euclidean distance with ClustVis.^27^ The microarray data have been deposited in Gene Expression Omnibus (GEO) under accession number GSE299842.

### Real-time quantitative PCR

*BCL2* expression was analyzed by real-time quantitative PCR (RT-qPCR) using TaqMan® probes (Hs00608023_m1, Thermo Fisher Scientific), using *GAPDH* (Hs99999905_m1) as endogenous control.^28^ For miRNA expression analysis, total RNA (10 ng) was reverse transcribed using the TaqMan® MicroRNA Reverse Transcription Kit (Thermo Fisher Scientific) and specific primers. RT-qPCR was performed with TaqMan® MicroRNA Assays: ID 002173 (hsa-miR-15b-3p), ID 002171 (hsa-miR-16-2-3p), and ID 001094 (RNU44) and ID 001006 (RNU48) as endogenous controls. All reactions were run on a 7900HT Fast Real-Time PCR System (Thermo Fisher Scientific). Relative expression levels were calculated using the ddCt method.

### Lentiviral constructs and transductions

Lentiviral (LV) constructs generation and LV particles production is detailed in Supplemental Data. ETPs were LV transduced for 24 h in RetroNectin® (Takara) pre-coated (50 μg/ml) 24-well plates and cultured (10^4^ cells per well) in RPMI 1640 with 10% FBS, rhSCF (100 IU/ml), rhIL-7 (200 IU/ml) (NIBSC) and rhFLT3L (50 ng/ml, PeproTech). LV transduction of patient-derived T-ALLs and cell lines was performed as described.^29^

### Flow cytometry, apoptosis and cell cycle analysis

Mouse anti-human mAbs used for cell surface staining are described in Supplemental Data. Apoptosis was assessed using the PE-Annexin-V Apoptosis Detection Kit I (BD Biosciences). Intracellular BCL-2 staining was performed with Cytofix/Cytoperm (BD Biosciences) and AlexaFluor-647-coupled anti-human BCL-2 (BD Biosciences). For cleaved Caspase 3 and CYCLIN D3 staining, Fixation/TruePhos Perm Buffers (Biolegend), and either PE-Cy7-coupled anti-cleaved Caspase 3 (Asp175) (Cell Signaling Technology) or PE-coupled anti-CYCLIN D3 (Biolegend) were used. To synchronize cells in the G2/M-phase, T-ALL cell lines were treated with Nocodazole (50 ng/ml, Sigma-Aldrich) for 16 h, then washed and cultured in fresh medium. Cell cycle was analyzed in cells fixed with 2% paraformaldehyde at room temperature for 20 min and stored overnight in 70% ethanol at -20°C. Fixed cells were washed with PBS and stained with DAPI, Dihydrochloride (1 μg /ml) in PBS plus 100 μg/ml RNase A (Sigma-Aldrich) and, when indicated, with PE-coupled mouse anti-Ki-67 Set (BD Biosciences). Flow cytometry was performed using a FACSCanto II and FlowJo v10 software (BD Biosciences).

### Western Blotting

Whole cell lysates prepared using RIPA buffer were separated by 10% sodium dodecyl sulfate–polyacrylamide gel electrophoresis (BioRad), transferred to polyvinylidene difluoride membranes (Merck), and incubated with Abs against BCL-2 (Cell Signaling Technology), CYCLIN D3 (BD Biosciences), and α-TUBULIN (Sigma-Aldrich). After washing, membranes were incubated with horseradish peroxidase–conjugated anti-mouse or anti-rabbit Abs (Jackson ImmunoResearch) for 1 h. Detection was done using Lumi-LightPLUS (Roche), and visualized with LAS 4000 mini (GE Healthcare) or x-ray film.

### Fetal thymic organ culture assays

All animal procedures were approved by the Comunidad de Madrid Animal Research Review Board (PROEX002621). For fetal thymic organ culture (FTOC) assays, E14.5 Swiss embryos thymic lobes depleted of thymocytes were cocultured with transduced human ETPs (10^4^ cells/lobe) for 2 days in hanging-drops, and transferred to nuclepore filters (Millipore) resting on gelfoam sponges as described.^30^

### Patient-derived T-ALL xenograft (PDX) and subcutaneous tumor assays

Patient-derived xenografts (PDX) were performed as described.^29^ Briefly, transduced human T-ALL cells (10^5^ T-ALL5 or 3 x 10^4^ T-ALL8 cells) were injected into the retro-orbital sinus of sublethally irradiated (1.5 Gy) 6- to 10-week-old NOD.Cg-*Prkdc^scid^Il2rg^tm1Wjl^*/SzJ (NSG; The Jackson Laboratory) immunodeficient mice. Mice were monitored weekly by body weight and flow cytometry to track T-ALL expansion in PB, and were euthanized when disease symptoms appeared. Human leukemic cells in the thymus, BM, spleen and liver were detected by flow cytometry. For subcutaneous tumor assays, 1-5 x 10^6^ transduced T-ALL cells were injected into the right flank of 6- to 10-week-old NSG mice. Tumor growth was measured every 2-3 days using a digital caliper, and mice were euthanized when tumor volume reached 3000 mm^3^.^29^

### Statistical analysis

A detailed description of the statistical analysis can be found in Supplemental Data.

## RESULTS

### Stage-specific regulation of miR-15b/16-2 expression along human T-cell development

We used a custom-designed microarray platform^24–26^ to analyze miRNA expression profiles across distinct stages of T-cell development in the human thymus (**Supplemental Table 1**). Our analysis revealed dynamic miRNA expression patterns (**Figure 1A**), indicating that specific miRNAs may contribute to defined developmental checkpoints. Notably, miR-16-2*, corresponding to mature miR-16-2-3p, was upregulated during ETP development, peaking at the DP CD3^-^ stage of resting thymocytes which lack TCRαβ expression and are positioned immediately downstream of pre-TCR-mediated β-selection.^31^ These findings were validated by RT-qPCR across sequential intrathymic stages. Since the *MIR16-2* gene is clustered with *MIR15B* within an intron of the *SMC4* gene (chromosome 3q26),^32^ we also analyzed mature miR-15b-3p expression. Both miR-15b-3p and miR-16-2-3p, showed a coordinated expression, which increased during early T-cell development, peaked at the DP CD3^-^ stage, coincident with pre-TCR downregulation after β-selection, and was markedly downregulated in more mature DP and SP thymocytes expressing TCRαβ (**Figure 1B**). The strong correlation between miR-15b-3p and miR-16-2-3p expression (**Figure 1C**) further supports the coordinated regulation of miR-15b and miR-16-2 members of the miR-15b/16-2 cluster during human T-cell development.

**Figure 1.**
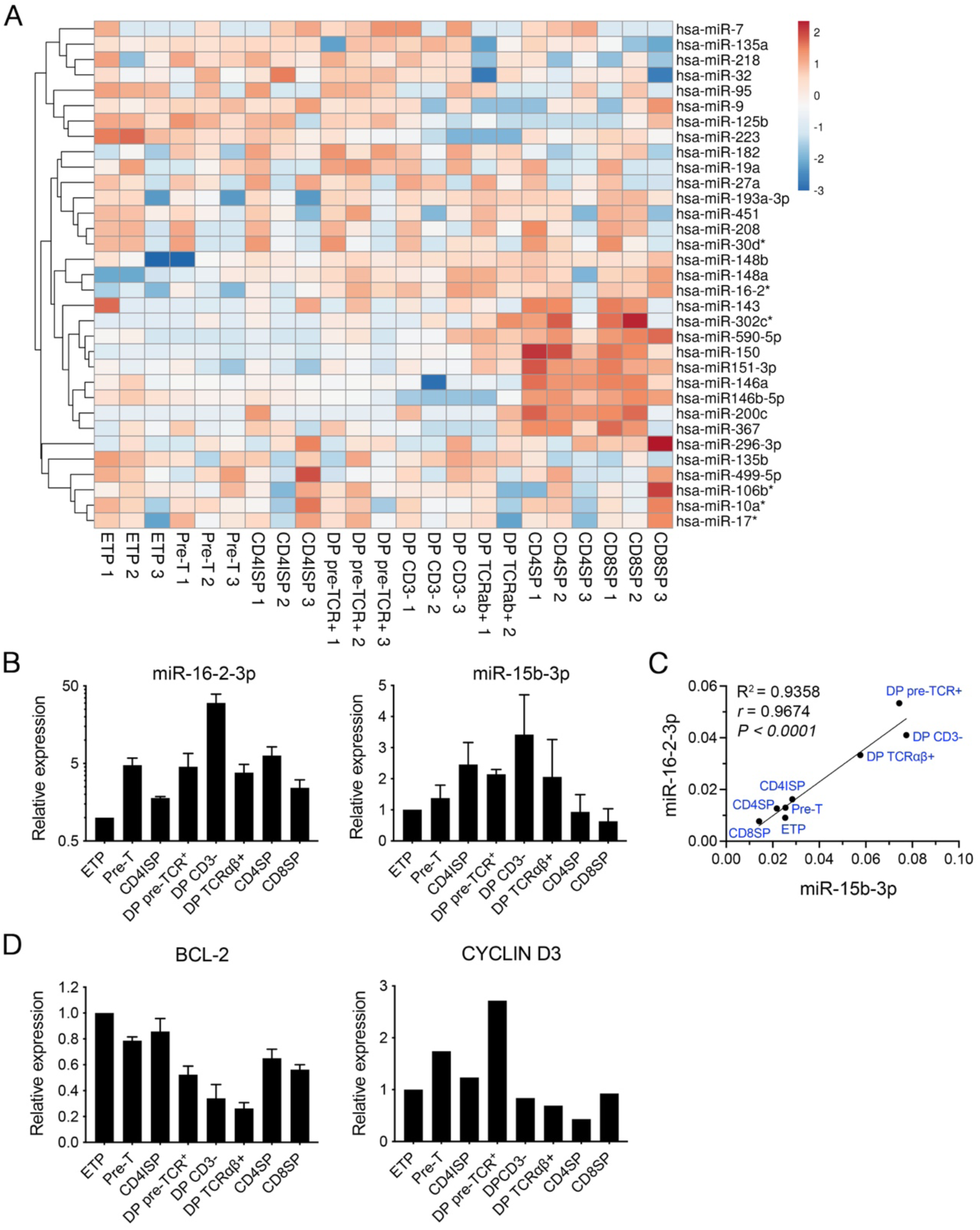
Stage-specific expression of the miR-15b/16-2 cluster in human intrathymic T-cell development. (A) Heatmap showing the hierarchical clustering of miRNAs expression in human intrathymic T-cell populations representative of ETP, Pre-T, CD4ISP, DP pre-TCR^+^, DP CD3^-^, DP TCRαβ^+^, CD4 SP and CD8 SP developmental stages. The color scale is based on z-score-scaled miRNA expression. The z-score distribution ranges from −3 (blue) to +2 (red). Each row represents a human miRNA, each column represents an intrathymic T-cell population, and 1-3 indicate three independent thymus samples. (B) Expression of miR-16-2-3p (hsa-miR-16-2*) (left) and miR-15b-3p (hsa-miR-15b*) (right) in human intrathymic T-cell populations shown in (A), determined by RT-qPCR. Results represent relative expression (RQ) compared to ETPs. Data are shown as mean values ± standard error of the mean (SEM) of three independent thymus samples run in triplicate. RNAU44/U46 were used as endogenous controls. (C) Correlation analysis between miR-15b-3p and miR-16-2-3p expression in the human intrathymic T-cell populations shown in (B). Expression levels were calculated by the ΔΔCT method using RNAU44/U46 as endogenous controls. Goodness-of-fit of linear regression (R^2^), Pearson correlation coefficient (*r)* and statistical significance (*P*) are indicated. (D) BCL-2 (left) and CYCLIN D3 (right) protein expression analyzed by intracellular flow cytometry in the human intrathymic T-cell populations shown in A. Data are geometric mean of fluorescence intensity (gMFI) of each intrathymic population relative to ETP values of 3 (BCL-2, ± SEM), and 2 (CYCLIN D3) independent thymus samples.

Known targets of the miR-15b/16-2 cluster include genes involved in cell survival and proliferation, such as *BCL2* and *CCND3* (CYCLIN D3).^33–35^ Consistent with this, increasing miR-15b and miR-16-2 expression from ETPs to β-selected DP CD3^-^ thymocytes was associated with decreased BCL-2 protein levels (**Figure 1D**). CYCLIN D3 also declined sharply at the DP CD3^-^ stage, coinciding with miR-15b-3p and miR-16-2-3p maximal expression (**Figure 1D**). These findings suggest that the miR-15b/16-2 cluster may regulate survival and/or proliferation at early stages of human T-cell development.

### Overexpression of the miR-15b/16-2 genomic cluster in human ETPs impairs expansion of the developing progeny in fetal thymic organ cultures

To directly investigate the role of miR-15b/16-2 in early human T-cell development, ETPs from human postnatal thymus were transduced with a bicistronic lentiviral vector encoding the human *MIR15B* and *MIR16-2* genes, along with GFP as a cell tracer (**Figure 2A**). The developmental potential of ETPs overexpressing miR-15b/16-2 was then assessed using a FTOC assay, in comparison with ETPs transduced with a control vector expressing only GFP (mock) (**Figure 2B**). Flow cytometry analyses revealed a significant decrease (up to 10-fold) in the percentages of developing thymocytes expressing miR-15b/16-2 along culture, compared to mock-transduced thymocytes, which remained stable (**Figure 2C**), suggesting a proliferative disadvantage of the former. Consistently, a significant reduction in the absolute numbers of miR-15b/16-2-expressing cells was observed, compared to the mock-transduced controls (**Figure 2D**), which expanded robustly, similar to non-transduced (GFP^-^) thymocytes (**Figure 2E**). These findings suggest that miR-15b/16-2 impairs survival and/or proliferation of the ETP progeny developing in FTOC.

**Figure 2.**
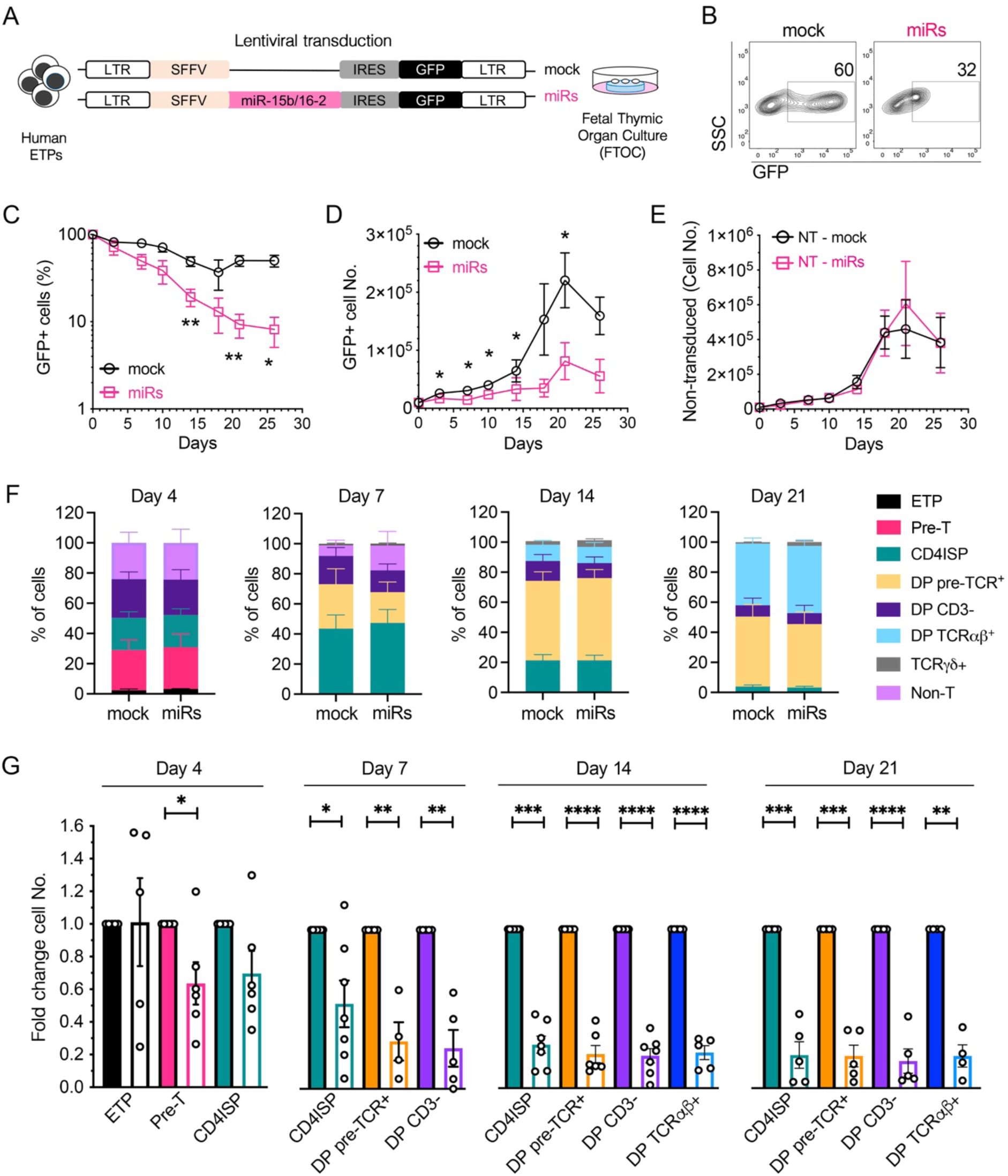
Overexpression of miR-15b/16-2 in human ETPs prevents the expansion of the ETP cell progeny. (A) Schematic representation of the lentiviral vectors used for ETP transduction, encoding either the miR-15b/16-2 cluster plus GFP as cell tracer (miRs) or GFP alone as control (mock). T-cell development of transduced ETPs was analyzed in a FTOC assay. (B) Flow cytometry analysis representative of the transduction efficiencies of human ETPs using the lentiviral vectors depicted in (A). Numbers indicate percentages of transduced (GFP^+^) cells. SSC, side-scatter. (C) Percentages of cells recovered at the indicated days from FTOCs seeded with mock- or miRs-transduced ETPs normalized to 100% of transduced cells at day 0. (D) Absolute numbers of mock- or miRs-transduced cells recovered from FTOCs in (C), normalized to an input of 10^4^ transduced ETPs. (E) Absolute numbers of non-transduced (NT) cells recovered from FTOCs in (C) seeded with mock- or miRs-transduced ETPs normalized to an input of 10^4^ non-transduced ETPs. (F) Percentages of cell populations generated from mock- or miRs-transduced ETPs at the indicated days of culture in FTOC. (G) Relative numbers of T-cell populations generated from miRs-transduced ETPs (empty bars) at the indicated days of culture in FTOC normalized to control populations derived from mock-transduced ETPs (filled bars). Data in (C-G) represent mean values ± SEM from at least 4 independent experiments. Statistical differences were determined by Student’s t tests with Holm-Šídák posttest analysis (C-D), Wilcoxon matched-pairs signed-rank test with False Discovery Rate (FDR) correction for multiple comparisons (E), two-way ANOVA with Holm-Šídák posttest analysis (F), and one-sample t test (G). \**P < .05*; \*\**P < .01*; \*\*\**P < .001*; \*\*\*\**P < .0001*.

To identify the specific developmental stage/s affected by miR-15b/16-2 overexpression, we analyzed the phenotype of progeny derived from either miR-15b/16-2- or mock-transduced ETPs in FTOC. No significant differences were observed in the distribution of T-cell subsets at various culture time points (**Figure 2F**), indicating that miR-15b/16-2 does not affect the differentiation potential of human ETPs in FTOC. However, the absolute number of cells in the miR-15b/16-2-expressing progeny was significantly reduced compared to mock-transduced cells (1.5- to 6-fold at different time points) (**Figure 2G**). This reduction affected all stages downstream of ETPs from pre-T to DP TCRαβ^+^ cells, resulting in up to an 80% decrease in developing T cells. In summary, overexpression of miR-15b/16-2 in human ETPs does not compromise their differentiation capacity, but significantly impairs expansion of the developing progeny at successive maturation stages in FTOC.

### miR-15b/16-2 overexpression impairs *in vitro* expansion and *in vivo* tumor progression of human T-ALL

Previous studies in mice have shown that the miR-15b/16-2 cluster acts as a B-cell tumor suppressor.^36^ To determine whether its inhibitory effect on cell expansion also applies to tumors derived from T-cell progenitors, we analyzed the *in vitro* proliferation of four human T-ALL cell lines (CUTLL1, JURKAT, HPBALL, SUPT1) after lentiviral transduction with miR-15b/16-2. This resulted in a 5- to 20-fold increase in miR-15b/16-2 expression compared to mock controls (**Supplemental Figure 1A**). All miR-15b/16-2-transduced cell lines showed a marked decrease in both relative (**Figure 3A**) and absolute cell numbers in culture (**Supplemental Figure 1B**), indicating that miR-15b/16-2 impairs T-ALL cell expansion *in vitro*.

**Figure 3.**
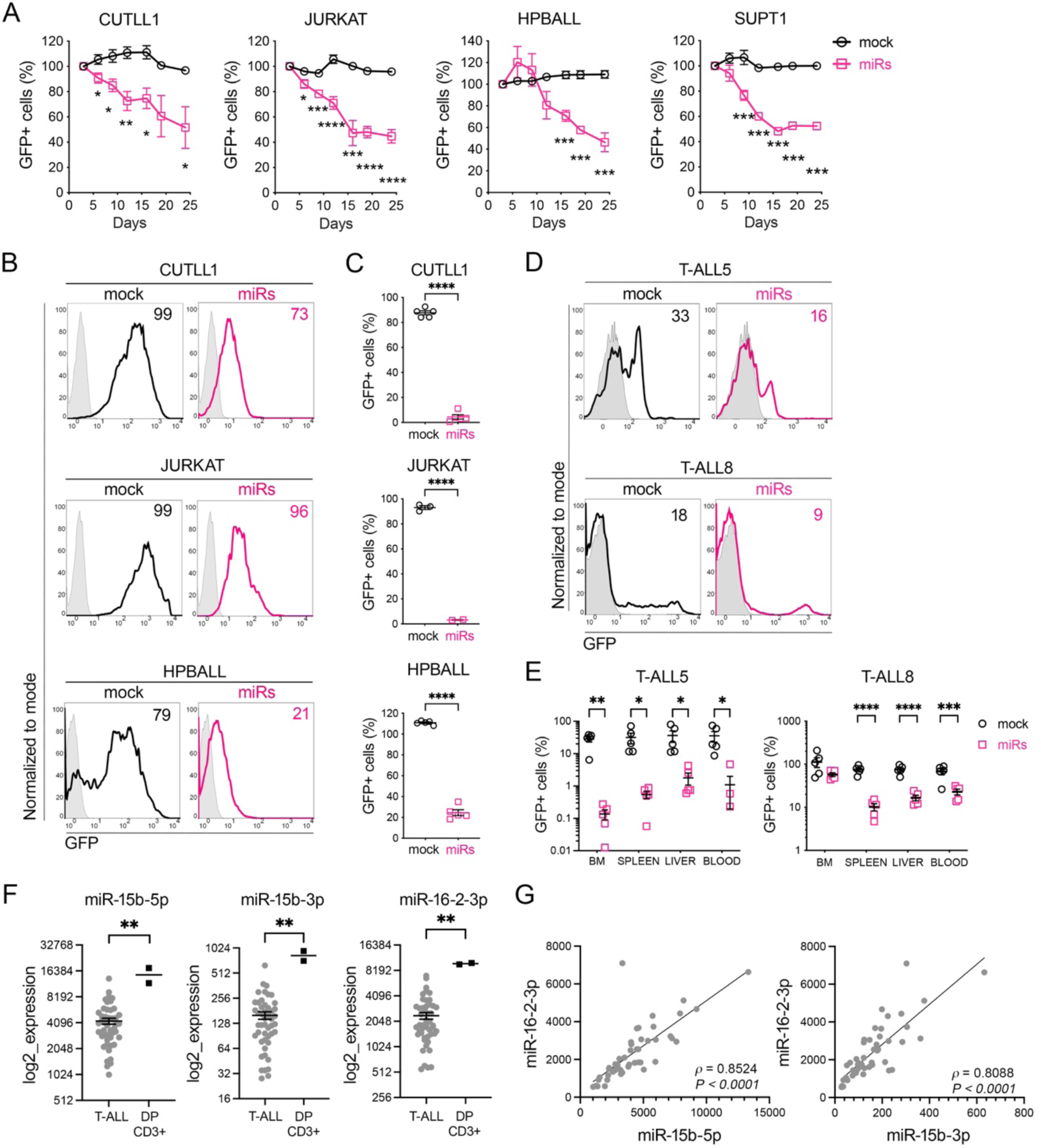
miR-15b/16-2 impairs *in vitro* cell expansion and *in vivo* tumor progression of T-ALL cells and is underexpressed in T-ALL patient samples. (A) *In vitro* cellular expansion of CUTLL1, JURKAT, HPBALL and SUPT1 human T-ALL cell lines lentivirally transduced with either miR-15b/16-2 plus GFP (miRs), or GFP alone (mock). Results are shown as mean percentages ± SEM of transduced (GFP^+^) cells recovered at the indicated days of culture normalized to 100% of transduced cells at day 0 (n ≥ 3). (B, C) *In vivo* progression of mock- and miRs-transduced CUTLL1, JURKAT, and HPBALL T-ALL cell lines subcutaneously transplanted into immunodeficient mice. Percentages of mock- and miRs-transduced cells determined by flow cytometry at the day of injection are shown in (B). Data in (C) show mean percentages of mock- and miRs-transduced T-ALL cells recovered from subcutaneous tumors, relative to percentages of transduced cells at the day of injection (CUTLL1, n = 5 mice/group; JURKAT, n ≥ 2 mice/group; HPBALL, n = 5 mice/group). (D, E) *In vivo* progression of mock- and miRs-transduced human primary T-ALL5 and T-ALL8 patient samples upon i.v. injection into immunodeficient mice. Percentages of mock- and miRs-transduced T-ALL5 and T-ALL8 cells determined by flow cytometry at the day of injection are shown in (D). Results in (E) show mean percentages ± SEM of mock- and miRs-transduced primary T-ALL5 and T-ALL8 cells recovered from the indicated organs of injected mice at 12.5-weeks or 6.5-weeks post-injection, respectively, relative to the percentage of transduced cells at the day of injection. (F) Decreased expression of miR-15b-5p (left), miR-15b-3p (middle) and miR-16-2-3p (right) in T-ALL patient samples (n = 48) compared to healthy DP CD3^+^ thymocyte samples (n = 2). Results are shown as mean values obtained from a publicly available small-RNA sequencing study (GSE89978).^13^ (G) Correlation analysis between miR-15b-5p and miR-16-2-3p (left), and miR-15b-3p and miR-16-2-3p (right) expression in T-ALL patient samples shown in (F). Spearman’s correlation coefficient (*π*) and statistical significance (*P*) are shown. Statistical differences were determined by Student’s t tests in A, C, E, and by Mann-Whitney in F. \**P < .05*; \*\**P < .01*; \*\*\**P < 0.001*; \*\*\*\**P < .0001*.

We next assessed the *in vivo* effects of miR-15b/16-2 overexpression using subcutaneous tumor assays in NSG mice with miR-15b/16-2- or mock-transduced T-ALL cells (CUTLL1, JURKAT and HPBALL; **Figure 3B**). While the proportions of mock-transduced cells remained stable in subcutaneous tumors (**Figure 3C**) compared with the day of injection (**Figure 3B**), miR-15b/16-2-transduced cells were dramatically reduced by up to 20-fold (CUTLL1), 30-fold (JURKAT), and 5-fold (HPBALL) (**Figure 3C**), indicating that miR-15b/16-2 impairs tumor progression *in vivo*. To confirm whether miR-15b/16-2 also impairs progression of primary T-ALL, we used a patient-derived xenograft (PDX) model in NSG mice with two T-ALL samples (T-ALL5 and T-ALL8). Mice received either mock- or miR-15b/16-2-transduced cells (**Figure 3D**) and were monitored every two days for body weight changes and weekly for disease progression in PB, and were euthanized when disease symptoms appeared. Strikingly, T-ALL cells infiltrating the blood, BM, spleen, and liver showed significantly reduced proportions of miR-15b/16-2- transduced cells, compared to input levels (100-1000-fold for T-ALL5; 5-50-fold for T-ALL8) (**Figure 3E**). In contrast, mock-transduced T-ALL counterparts remained stable in all organs analyzed. Moreover, absolute numbers of miR-15b/16-2-transduced T-ALL cells were also significantly reduced compared with mock-transduced controls (**Supplemental Figure 2A**), which expanded similarly to non-transduced cells (**Supplemental Figure 2B**). This suggests that ectopic miR-15b/16-2 expression hinders *in vivo* cell expansion and tumor progression of patient T-ALL, supporting its tumor suppressor role.

To assess the clinical relevance of our findings, we analyzed publicly available data from Wallaert et al. (GSE89978),^13^ which profiled miRNA expression in a cohort of 48 T-ALL patient samples. This study showed that expression of three mature miRNAs from the miR-15b/16-2 genetic cluster (*i.e.* miR-15b-5p, miR-15b-3p and miR-16-2-3p) was significantly downregulated in T-ALL samples versus healthy DP CD3^+^ thymocytes (**Figure 3F**), and confirmed a strong correlation between miR-15b and miR-16-2 expression (**Figure 3G**). These findings further support a tumor suppressor function for miR-15b/16-2 in T-ALL.

### miR-15b/16-2 induces T-ALL cell apoptosis partly through BCL-2 downregulation

The reduction in T-ALL cell expansion observed upon miR-15b/16-2 overexpression *in vitro* and *in vivo* appears to result from increased apoptosis and/or reduced proliferation. Flow cytometry showed that miR-15b/16-2 increased early (Annexin-V^+^ 7-AAD^-^) and late apoptosis (Annexin-V^+^ 7-AAD^+^) of T-ALL cell lines (**Figure 4A**), resulting in a 2- to 7-fold increase in total apoptotic cells compared to mock-transduced controls (**Figure 4B**). miR-15b/16-2-dependent apoptosis was further confirmed by cleaved caspase 3 intracellular staining (**Supplemental Figure 3A, B**). Since BCL-2 is a known target of the miR-15/16 family^33^ and a key antiapoptotic factor,^37^ we next examined whether miR-15b/16-2 induces apoptosis via BCL-2 downregulation. Both western blot (**Figure 4C, D**) and flow cytometry assays (**Supplemental Figure 3C, D**) confirmed a significant reduction of BCL-2 in miR-15b/16-2-transduced compared with mock-transduced T-ALL cell lines. Similar effects were observed in patient-derived T-ALL cells (T-ALL5 and T-ALL8), where ectopic expression of miR-15b/16-2 led to increased apoptosis both *in vitro* (**Figure 4E, F, and Supplemental Figure 3E, F**) and *in vivo* (**Supplemental Figure 3G**), and to BCL-2 downregulation compared with mock controls (**Figure 4G, H**).

**Figure 4.**
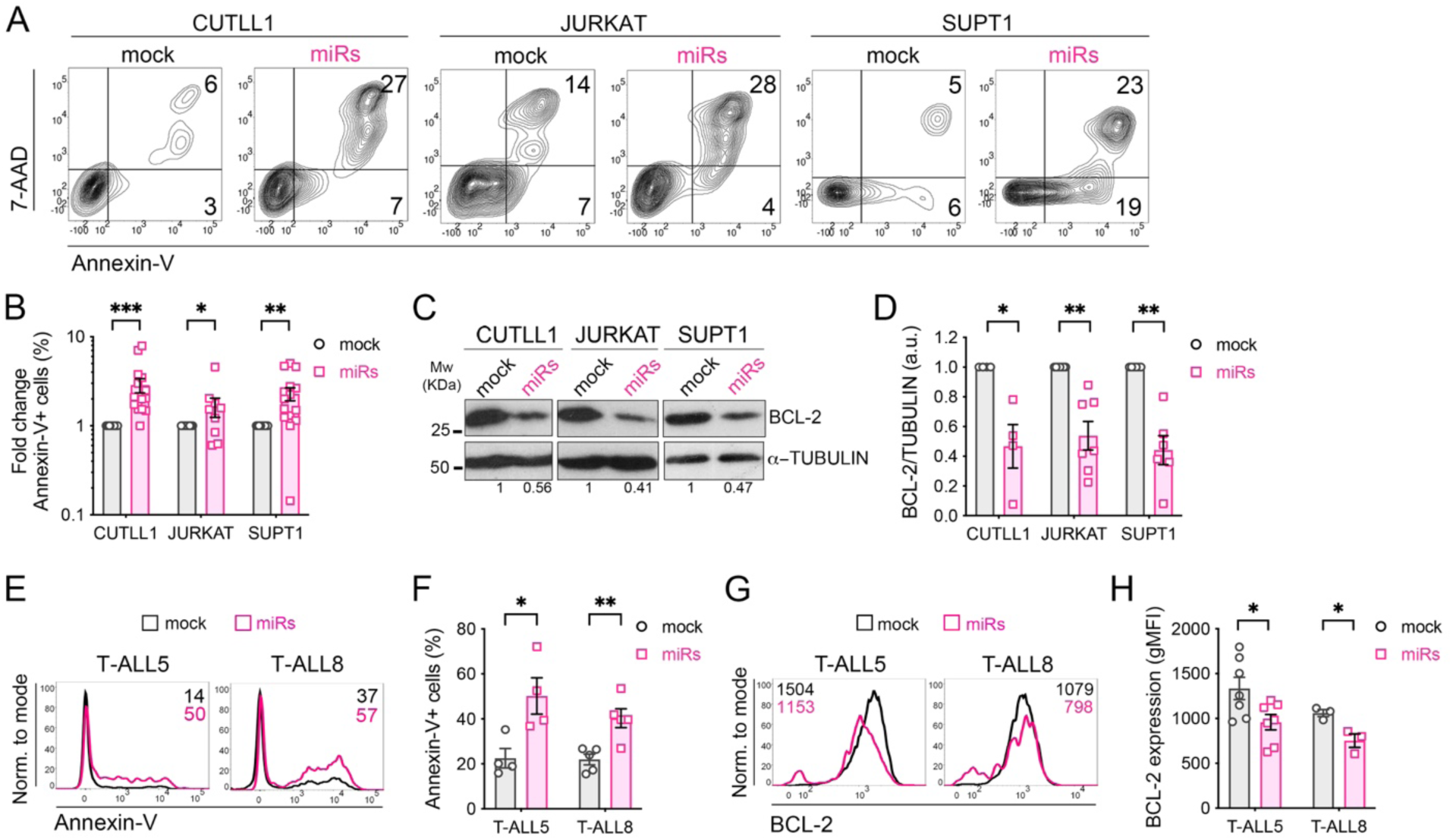
Ectopic expression of miR-15b/16-2 induces cell apoptosis and BCL-2 downregulation in human T-ALL. (A) Representative dot plots of apoptosis detected by Annexin-V/7-AAD labeling in CUTLL1, JURKAT and SUPT1 human T-ALL cell lines lentivirally transduced with either miR-15b/16-2 plus GFP (miRs) or GFP alone (mock) as control. Numbers indicate percentages of positive cells in each quadrant at day 3 post-transduction. (B) Quantification of apoptotic (Annexin-V^+^) miRs-transduced T-ALL cells relative to apoptotic mock-transduced T-ALL cells. Data are shown as mean values ± SEM of n ≥ 8 independent experiments (CUTLL1, n = 15; JURKAT, n = 8; SUPT1, n = 15). Statistical differences were determined by Wilcoxon signed-rank test (CUTLL1, JURKAT), and one-sample t test (SUPT1). (C) Representative immunoblot of BCL-2 expression in the indicated mock- and miRs-transduced human T-ALL cell lines. α-TUBULIN was used as protein loading control. Molecular weight (Mw) is indicated on the left. BCL-2 / α-TUBULIN expression ratio in miRs-transduced relative to mock-transduced T-ALL cell lines shown. (D) Quantification of BCL-2 / α-TUBULIN expression ratio in different experiments performed as in (C). Graphs represent mean values ± SEM of n ≥ 4 independent experiments (CUTLL1, n = 4; JURKAT, n = 7; SUPT1, n = 6). Statistical differences were determined by one-sample t test; a.u., arbitrary units. (E) Representative flow cytometry histograms of apoptosis levels determined by Annexin-V staining in human primary T-ALL5 and T-ALL8 patient samples transduced with either miR-15b/16-2 plus GFP (miRs) or GFP alone (mock). Numbers indicate percentages of apoptotic cells at day 3 post-transduction. (F) Mean percentages ± SEM of apoptotic (Annexin-V^+^) miRs- and mock-transduced T-ALL patient cells analyzed in independent experiments (T-ALL5, n = 4; T-ALL8, n = 5). Statistical differences between groups were determined by Student’s t test. (G) Representative flow cytometry histograms of BCL-2 protein expression in mock- and miRs-transduced human primary T-ALL5 and T-ALL8 patient samples at day 3 post-transduction. Numbers indicate gMFI of BCL-2 expression. (H) Quantification of BCL-2 expression levels in mock- and miRs-transduced human primary T-ALL patient samples analyzed as in (G). Data are shown as mean values ± SEM of independent experiments (T-ALL5, n = 7; T-ALL8, n = 3). Statistical differences were determined by Student’s t tests. \**P < .05;* \*\**P < .01*; \*\*\**P < .001*.

To directly investigate whether miR-15b/16-2-induced downregulation of BCL-2 leads to apoptosis in T-ALL cells, we engineered a BCL-2 rescue vector encoding a miR-15/16-resistant BCL-2 together with miR-15b/16-2 (**Supplemental Figure 4A**), which induces a significant increase of BCL-2 expression in transduced cells (**Supplemental Figure 4B-D**). We found that BCL-2 overexpression only partially rescued apoptosis induced by miR-15b/16-2 (**Figure 5A, B**), and did not restore cell expansion *in vitro* (**Figure 5C**) or *in vivo*, as subcutaneous tumor assays revealed no differences in the proportions of JURKAT cells transduced with either miR-15b/16-2 or the rescue vector (**Figure 5D, E**). More importantly, equivalent results were obtained with patient-derived T-ALL cells (T-ALL8) in a PDX model showing that tumor progression in the BM, spleen and liver was similar in NSG mice infused with T-ALL cells transduced with the BCL-2-rescue vector or with miR-15b/16-2 (**Figure 5F, G**), despite significant BCL-2 overexpression was observed in the former (**Figure 5H**). In summary, miR-15b/16-2 induces apoptosis in T-ALL cells partly through BCL-2 downregulation, but other miR-15b/16-2-dependent mechanisms likely contribute to the impaired T-ALL tumor growth *in vivo*.

**Figure 5.**
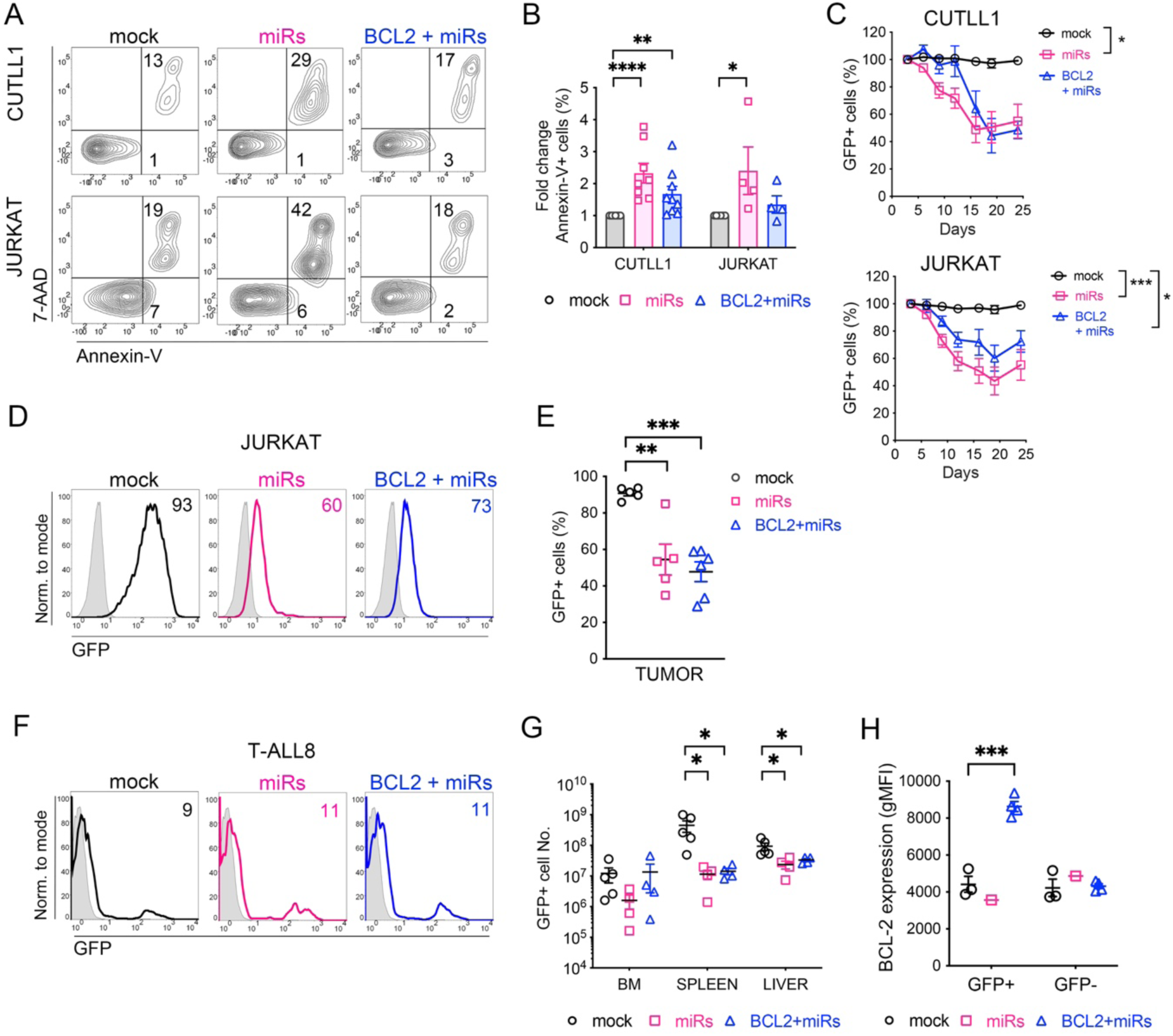
Ectopic expression of BCL-2 partially rescues miR-15b/16-2-induced apoptosis but is not sufficient to rescue T-ALL tumor expansion. (A) Representative flow cytometry analysis of apoptosis levels detected by Annexin-V/7-AAD labeling in CUTLL1 and JURKAT human T-ALL cell lines lentivirally transduced with either miR-15b/16-2 plus GFP (miRs), or a BCL-2 rescue vector encoding BCL-2, miR-15b/16-2 and GFP (BCL2 + miRs), or GFP alone (mock) as control. Numbers indicate the percentages of positive cells in each quadrant at day 3 post-transduction. (B) Quantification of apoptotic (Annexin-V^+^) cells among miRs-transduced or BCL2 + miRs-transduced T-ALL cell lines relative to apoptotic mock-transduced T-ALL cells. Data are shown as mean values ± SEM of independent experiments (JURKAT, n = 4; CUTLL1, n ≥ 9). Statistical differences were determined by Kruskal-Wallis with Dunn posttest analysis. (C) *In vitro* cellular expansion of CUTLL1 and JURKAT human T-ALL cell lines lentivirally transduced as in (A). Results are shown as mean percentages ± SEM of transduced (GFP^+^) cells recovered at the indicated days of culture normalized to 100% of transduced cells at day 0 (n ≥ 4). Statistical differences were determined by one-way ANOVA with Tukey posttest analysis. (D, E) *In vivo* progression of JURKAT T-ALL cells transduced with mock-, miRs- and BCL2 + miRs-encoding lentiviral vectors and subcutaneously transplanted into immunodeficient mice. Transduction efficiencies determined by flow cytometry at the day of injection are shown in (D). Data in (E) are mean percentages ± SEM of JURKAT cells transduced with the indicated vectors recovered from subcutaneous tumors of transplanted mice relative to the percentage of transduced cells at the day of injection (n ≥ 5). Statistical differences were determined by one-way ANOVA with Tukey posttest analysis. (F, G) *In vivo* progression of mock-, miRs- and BCL2 + miRs-transduced primary T-ALL8 patient cells i.v.-injected into immunodeficient mice. Percentages of transduced T-ALL8 cells determined by flow cytometry at the day of injection are shown in (F). Results in (G) show mean absolute numbers ± SEM of mock-, miRs- and BCL2 + miRs-transduced primary T-ALL8 cells recovered from the indicated organs of injected mice at 6.5-weeks post-injection normalized to 10^5^ injected transduced cells. Statistical differences were determined by one-way ANOVA with Tukey posttest analysis. (H) BCL-2 protein expression determined by flow cytometry in either non-transduced (GFP^-^) or transduced T-ALL8 cells recovered from the BM of transplanted immunodeficient mice in (G) at 6.5-weeks post-transplant. Data are shown as gMFI of BCL-2 ± SEM. Statistical differences were determined by one-way ANOVA with Tukey posttest analysis. \**P < .05*; \*\**P < .01;* \*\*\**P < .001;* \*\*\*\**P < .0001*.

### miR-15b/16-2 overexpression induces CYCLIN D3 downregulation and impairs cell cycle progression in T-ALL

Cell cycle progression is partly regulated by several cyclins, including CYCLIN D3, which are direct targets of the miR-15/16 family.^34,36,38–41^ To assess the impact of miR-15b/16-2 overexpression in T-ALL, we analyzed cell cycle status by DAPI and Ki-67 staining of T-ALL cell lines transduced with miR-15b/16-2. Flow cytometry showed that T-ALL cells overexpressing miR-15b/16-2 accumulated in the G0 phase compared to mock-transduced controls (**Figure 6A, B**). To further investigate cell cycle disruption induced by miR-15b/16-2, we synchronized cells using Nocodazole to induce G2/M arrest. Flow cytometry revealed that miR-15b/16-2 overexpression impaired G2/M arrest in all T-ALL cell lines, and increased G0/G1 accumulation compared to mock-transduced controls (**Figure 6C, D**), indicating delayed progression from G0/G1 to S and G2/M phases. In addition, after Nocodazole release, miR-15b/16-2-expressing T-ALL cells showed delayed re-entry into G0/G1 and prolonged S/G2/M arrest (**Figure 6E-G**). Together, these findings indicate that miR-15b/16-2 disrupts normal cell cycle progression, contributing to impaired T-ALL proliferation.

**Figure 6.**
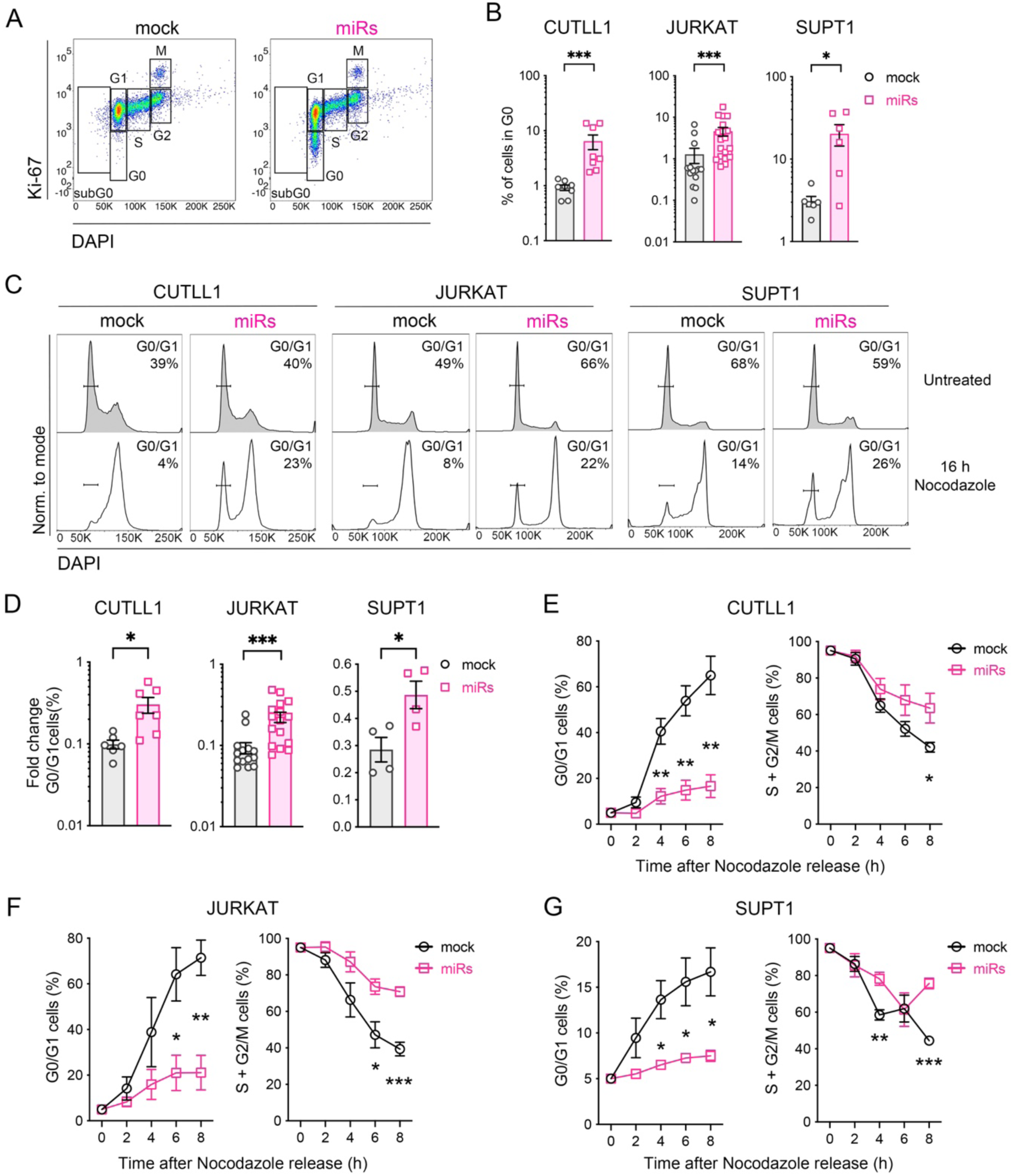
miR-15b/-16-2 overexpression prevents cell cycle progression in T-ALL. (A) Representative flow cytometry of cell cycle distribution analyzed by Ki-67/DAPI staining in CUTLL1 T-ALL cells lentivirally transduced with miR-15b/16-2 plus GFP (miRs) or with GFP alone (mock). Gates show cells in sub-G0, G0, G1, S, M and G2 cell cycle phases. (B) Percentages of cells in G0 in mock- or miRs-transduced CUTLL1, JURKAT and SUPT1 human T-ALL cell lines analyzed as in (A). Data are mean percentages ± SEM of independent experiments (CUTLL1, n ≥ 8; JURKAT; n ≥ 14 SUPT1, n ≥ 4). Statistical differences were determined by Student’s t test (SUPT1), or Mann-Whitney’s test (CUTLL1, JURKAT). (C) Representative cell cycle distribution in DAPI-stained mock- or miRs-transduced CUTLL1, JURKAT and SUPT1 T human T-ALL cells analyzed by flow cytometry in Nocodazole-based synchronization assays. Numbers indicate percentages of cells in G0/G1 before (grey histograms), and after (empty histograms) 16 h of Nocodazole treatment to induce G2/M cell cycle arrest. (D) Fold change of cell percentages in G0/G1 after 16 h of Nocodazole treatment relative to G0/G1 percentages of untreated cells in mock- and miRs-transduced T-ALL cell lines assayed as in (C). Data are mean values ± SEM of independent experiments (CUTLL1, n ≥ 6; JURKAT, n ≥ 14; SUPT1, n = 4). Statistical differences were determined by Student’s t test (CUTLL1, SUPT1), or Mann-Whitney’s test (JURKAT). (E-G) Percentages of cells in G0/G1 (left) and S+G2/M (right) in mock- and miRs-transduced CUTLL1 (E), JURKAT (F) and SUPT1 (G) T-ALL cell lines, at different time points after Nocodazole washout. Data were normalized to 5% of cells in G0/G1 and 95% of cells in S+G2/M after 16 h Nocodazole-treatment. Data are mean values ± SEM of independent experiments (CUTLL1, n = 4; JURKAT, n = 4; SUPT1, n = 4). Statistical differences were determined by Student’s t test with Holm-Šídák’s posttest for multiple comparisons at different times. \**P < .05;* \*\**P < .01; ***P < 0.001; **** P < .0001*.

These results prompted us to examine expression of CYCLIN D3, a known miR-15/16 target,^34,35,42,43^ implicated in T-cell development and T-ALL pathogenesis.^44^ Western blot and flow cytometry analyses showed significantly reduced CYCLIN D3 levels in miR-15b/16-2-transduced CUTLL1 and JURKAT cells, compared with mock-transduced controls (**Figure 7A, B; Supplemental Figure 5A, B**). Cells with reduced CYCLIN D3 levels were predominantly arrested at G0/G1 (**Figure 7C, D**), suggesting that miR-15b/16-2 induces G0/G1 arrest via CYCLIN D3 suppression. Notably, miR-15b/16-2 overexpression also induced CYCLIN D3 downregulation in primary T-ALL patient cells (**Figure 7E, F**).

**Figure 7.**
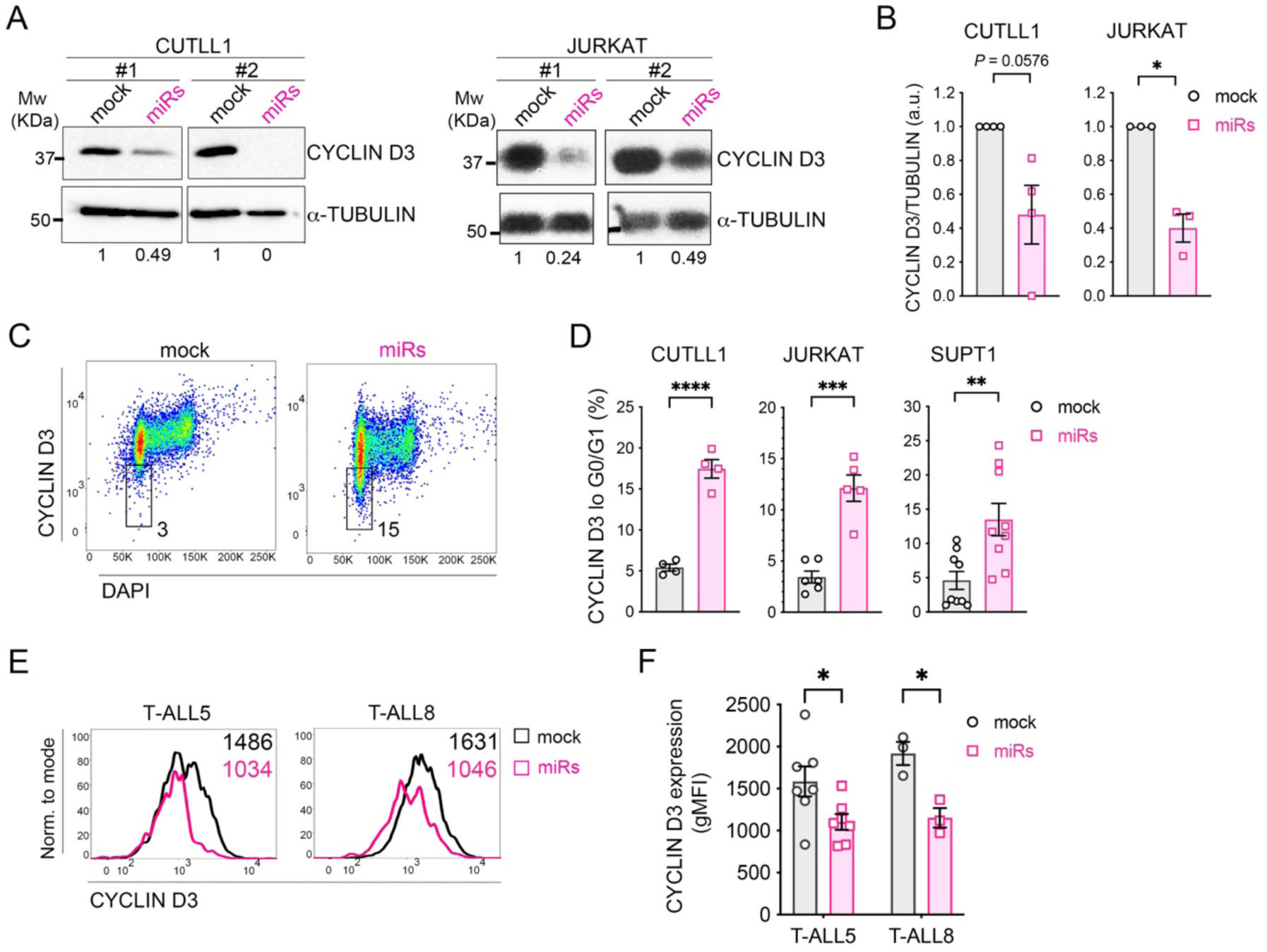
miR-15b/16-2 represses the expression of CYCLIN D3 in human T-ALL. (A) Representative immunoblots of CYCLIN D3 expression in CUTLL1 and JURKAT T-ALL cells transduced with miR-15b/16-2 plus GFP (miRs) or with GFP alone (mock). α-TUBULIN was used as protein loading control. Molecular weight (Mw) in KDa is indicated on the left. #1 and #2 represent two independent experiments. CYCLIN D3 / α-TUBULIN expression ratio in miRs-transduced relative to mock-transduced T-ALL cells is shown. (B) Quantification of CYCLIN D3 / α-TUBULIN expression ratio in different experiments performed as in (A). Graphs represent mean values ± SEM of independent experiments (CUTLL1, n = 4; JURKAT, n = 3). Statistical differences were determined by one-sample t test; a.u., arbitrary units. (C) Representative flow cytometry analysis of CYCLIN D3 expression at distinct cell cycle phases determined by DAPI staining of mock- and miRs-transduced JURKAT cells at day 7 post-transduction. Numbers indicate percentages of cells at G0/G1 expressing low levels of CYCLIN D3. (D) Mean percentages ± SEM of cells in G0/G1 expressing low CYCLIN D3 levels (CYCLIN D3^lo^) in the indicated mock- and miRs-transduced T-ALL cell lines analyzed as in (C). Statistical differences between groups were determined by Student’s t test at day 7 post-transduction (CUTLL1, n = 4; JURKAT, n = 5), or by Mann-Whitney’s test at day 4 post-transduction (SUPT1, n = 9). (E) Representative flow cytometry histograms of CYCLIN D3 protein expression in mock- and miRs-transduced human primary TALL-5 and T-ALL8 patient samples at day 4 post-transduction. Numbers indicate gMFI of CYCLIN D3 expression. (F) Quantification of CYCLIN D3 expression levels in mock- and miRs- transduced human primary T-ALL patient samples analyzed as in (E). Data are mean gMFI ± SEM of independent experiments (T-ALL5, n = 7; T-ALL8, n = 3). Statistical differences were determined by Student’s t test. \**P < .05*; \*\**P < .01;* \*\*\**P < .001;* \*\*\*\**P < .0001*.

## DISCUSSION

In this study, we conducted a microarray analysis to investigate miRNA expression across human T-cell development, aiming to identify stage-specific miRNA signatures, as indicators of functional involvement in particular differentiation processes. Our goal was to uncover potential links between individual miRNA functions, their physiological targets during T lymphopoiesis, and their potential impact on T-ALL initiation and progression. The analysis revealed a characteristic expression pattern for miR-16-2-3p, which increases in early proliferating T-cell progenitors, peaks in β-selected DP CD3^-^ resting thymocytes, and then declines in more mature DP and SP TCRαβ^+^ thymocytes. A coordinated expression of miR-15b-3p was also observed, highlighting the miR-15b/16-2 cluster as a key regulator of progenitor expansion during early T-cell development. Supporting this, overexpression of miR-15b/16-2 in ETPs impaired progeny expansion, leading to defective production of mature TCRαβ^+^ thymocytes. We conclude that miR-15b/16-2 fine-tunes gene expression networks that first enable early progenitor expansion and later limit proliferation following successful β-selection. This regulatory role appears crucial in the context of T-ALL pathogenesis, as miR-15b/16-2 overexpression impaired *in vivo* T-ALL progression, while its downregulation is frequently observed across T-ALL subtypes.^13^ Overall, our findings uncover a novel tumor suppressor function for the miR-15b/16-2 cluster in T-ALL.

The proposed role of miR-15b/16-2 in early T-cell development appears similar to its recently reported function in the pre-B to immature B cell transition during B lymphopoiesis.^34^ Like in immature B cells, miR-15b/16-2 may suppress proliferation to enable differentiation of resting thymocytes after β-selection. This conserved function reflects the shared need for sequential IL-7R and pre-TCR (or pre-BCR) signaling in T- and B-cell progenitor expansion, respectively.^28,45,46^ Therefore, miR-15b/16-2 may modulate IL-7R and pre-TCR signaling networks during T-cell development, similar to its role in IL-7R and pre-BCR signaling during B lymphopoiesis. This regulation may be essential for preventing early lymphoid malignancies, such as T- and B-ALL leukemias, which often result from uncontrolled proliferation in early lymphopoiesis.^47–49^ Since both IL-7R and pre-TCR critically contribute to T-ALL development and progression,^23,50–52^ miR-15b/16-2 likely participates in a regulatory loop with these pathways, as seen in B-cell development, thereby contributing to T-ALL pathogenesis. However, the exact contribution of miR-15b/16-2 dysregulation to T-ALL remains to be established.

Identifying miR-15b/16-2 cluster as a tumor suppressor in T-ALL is consistent with the known roles of other miR-15/16 family members in hematological cancers, such as B-cell lymphoma,^36^ chronic lymphocytic leukemia (CLL),^53^ and acute myeloid leukemia.^41^ Mechanistically, miR-15b/16-2 may exert its T-ALL tumor-suppressive effects by promoting apoptosis and disrupting the cell cycle, primarily through repression of *BCL2* and *CCND3*, two key miR-15b/16-2 target genes in T-ALL pathogenesis^44,54–60^ that support IL-7R-mediated survival and proliferation,^61^ and pre-TCR-induced expansion.^44,62^ Nonetheless, since BCL-2 appears partially responsible of miR-15b/16-2-mediated apoptosis, other antiapoptotic miR-15b/16-2 targets like MCL-1 and BMI-1^63,64^ may also be involved. The observed cell cycle arrest in G0 and delayed G2/M progression is likely due to reduced CYCLIN D3 expression induced by miR-15b/16-2, as observed in other cell types.^65–67^Additionally, miR-15b/16-2 may impair exit from drug-induced G2/M arrest by downregulating regulators such as *Cdc27* and *Anapc1*^68^, essential components of the APC/C complex, whose inhibition leads to sustained Cdk1 activity and Cyclin B accumulation.^69–71^ Thus, CYCLIN D3-driven cell cycle disruption may be a key mechanism behind the tumor suppressor role of miR-15b/16-2 in T-ALL.

Finally, our findings showing downregulation of the miR-15b/16-2 cluster in a previously published cohort of T-ALL patient samples^13^ support the clinical relevance of our results, highlighting miR-15b/16-2 as potential therapeutic targets. This may be particularly important for BCL-2 inhibitor-resistant T-ALLs,^54,72^ where miR-15b/16-2-mediated suppression of cell cycle progression could offer new therapeutic opportunities. Recent advances in miRNA *in vivo* delivery systems using nanoparticle encapsulation^73^ have improved the feasibility of miRNA-based therapies. Notably, human-ferritin-based nanoparticles have recently been used *in vitro* to deliver miR-15a/16-1 to ferritin-receptor positive CLL cells.^74^ In conclusion, our results open new avenues for developing future therapies involving the encapsulation of miR-15b/16-2 in antibody- or aptamer-coated nanoparticles,^75–77^ designed to selectively and safely target malignant T cells via T-ALL-specific antigens, while sparing healthy T cells.

## ACKNOWLEDGEMENTS

The authors thank Eloísa Castillo for technical assistance. We thank the expert personnel within the Animal Facility and the Flow Cytometry Core at the Centro de Biología Molecular Severo Ochoa, and the Pediatric Cardiosurgery Units from Hospital Universitario La Paz and Hospital Universitario 12 de Octubre (Madrid, Spain) for postnatal thymus samples. This research was supported in part by grants PID2019-105623RB-I00, and PID2022-138880OB-I00 funded by Ministerio de Ciencia, Innovación y Universidades. Agencia Estatal de Investigación (MCIN/AEI/ 10.13039/501100011033) / European Regional Development Fund, European Union, and by grants from Fundación Unoentrecienmil and Fundación Inocente Inocente. Institutional grants from the Fundación Ramón Areces and Banco de Santander to the Centro de Biología Molecular Severo Ochoa are also acknowledged.

## AUTHORSHIP

S.G-G., designed and performed experiments, analyzed and interpreted data, and wrote the manuscript; M.J.G-L. design and performed experiments; M.G-P., P.F., and E.M-G. performed experiments; J.A, assisted with *in vivo* experiments; R.G. and C.M.C., designed and performed microarray assays, and interpreted microarray data; M.L.T. supervised the study and wrote the manuscript.

Conflict-of-interest disclosure: The authors declare no competing financial interests.

