## Supplemental Data for "A tumor suppressor role of the miR-15b/16-2 cluster in T-cell acute lymphoblastic leukemia"

### SUPPLEMENTAL METHODS

#### Lentiviral constructs

Lentiviral (LV) constructs encoding the human *MIR15B* (NR_029663.1, miRbase MI0000438) and *MIR16-2* (NR_029525.1, miRBase MI0000115) genomic cluster (Chr.3, NC_000003.12, GRCh38.p14) were generated by PCR amplification with the following primers: forward 5'AGATCTTCTATCACATAAGTGGAAAGAAAAAGACATTTAAG 3' and reverse 5'AGATCTGAATTCCTAGGTGCTTAGGTAAATCAAACACCAAGTGTA 3', and cloned upstream of an IRES-GFP cassette in the pHRSIN LV vector using BamHI/BglII and EcoRI (pHRSIN-miR-15b/16-2-IRES-GFP, miRs). The BCL-2 rescue vector was generated by cloning human *BCL2* CDS (NM_000633.2) upstream of miR-15b/16-2 in the pHRSIN-miR-15b/16-2-IRES-GFP LV vector using AscI and the following primers: forward 5' CCAGATCTGGCGCGCCCGCCACCATGGCGCACGCTGGGAGAACA 3' and reverse 5' CCAGATCTGGCGCGCCTCACTTGTGGCCCAGATAGGCACC 3' (pHRSIN-BCL2-miR-15b/16-2-IRES-GFP, BCL2+miRs). A GFP-alone LV vector was used as control (pHRSIN-IRES-GFP, mock). For LV supernatant production, HEK293T cells were transfected with each LV plasmid, gag-pol (psPAX2), and either VSV-G envelope (pMD2G) for T-ALL cell transduction, or MMLV envelope for ETPs, using JetPei (PolyPlus), and cultured for 48-72 hours.

#### Flow cytometry analysis

Mouse anti-human mAbs used for cell surface staining included: CD1a-PE (SFCI19Thy1A8), CD4-PE (SFCI12T4D11), CD4-PE-Cy5 (13B8.2), CD34-APC (581), TCRαβ-PE (IP26), TCRαβ-PE-Cy5 (IP26), TCRγδ-PE-Cy5 (5A6.E9) (Beckman Coulter); CD1a-APC (HI149), CD3-APC (UCHT1), CD3-PE (UCHT1), CD7-BV421 (M-T701), CD8-PE-Cy7 (RPA-T8), CD13-PE (WM15),

CD45-V450 (2D1) (BD Biosciences), and TCRαβ-biotin (IP26) and CD1a-biotin (HI149) (Biolegend). Biotinylated antibodies were developed using Brilliant Violet 421 Streptavidin (Biolegend). Background fluorescence was determined with irrelevant isotype-matched Abs (BD Biosciences).

**Statistical analysis**

Statistical analyses were conducted using GraphPad Prism (v9.5.0, GraphPad Software, San Diego, CA, USA). Normality was assessed with Shapiro-Wilk test. Comparisons between two groups were performed using a two-tailed Student's t-test for normally distributed data, or Mann-Whitney test for non-normally distributed data. To correct for multiple comparisons across time points, the Holm-Šídák method was applied, and adjusted *P*-values are reported. For comparisons among three groups, one-way ANOVA followed by Tukey's post hoc test was applied for normally distributed data, and Kruskal-Wallis test with Dunn's post hoc correction was used for non-normally distributed data. A one-sample t-test, or Wilcoxon signed-rank test, was used for comparisons between two groups when GFP-control transduced cells were normalized to 1 or 100. Data in graphs represent mean ± standard error of the mean (SEM). Simple linear regression analysis was used to assess the relationship between miR-15b-3p and 16-2-p expression. The strength and direction of the association were evaluated using Pearson's ($r$), or Spearman's ($\rho$), correlation coefficients. A significance level of $\alpha = 0.05$ was used for all statistical tests.

**SUPPLEMENTAL TABLE**

| **Population** | ***Phenotype*** |
|---|---|
| ETP | $CD34^{hi}$ $CD1a^{-}$ $CD4^{-}$ $CD8^{-}$ $CD3^{-}$ $TCR\alpha\beta^{-}$ |
| Pre-T | $CD34^{+}$ $CD1a^{+}$ $CD4^{-}$ $CD8^{-}$ $CD3^{-}$ $TCR\alpha\beta^{-}$ |
| CD4ISP | $CD4^{+}$ $CD1a^{+}$ $CD8^{-}$ $CD3^{-}$ $TCR\alpha\beta^{-}$ |
| DP pre-$TCR^{+}$ | $CD4^{+}$ $CD8^{+}$ $CD3^{lo}$ $TCR\alpha\beta^{-}$ |
| DP $CD3^{-}$ | $CD4^{+}$ $CD8^{+}$ $CD3^{-}$ $TCR\alpha\beta^{-}$ |
| DP $TCR\alpha\beta^{+}$ | $CD4^{+}$ $CD8^{+}$ $CD3^{+}$ $TCR\alpha\beta^{+}$ |
| CD4SP | $CD4^{+}$ $CD8^{-}$ $CD3^{+}$ $TCR\alpha\beta^{+}$ |
| CD8SP | $CD4^{-}$ $CD8^{+}$ $CD3^{+}$ $TCR\alpha\beta^{+}$ |

**Supplemental Table 1.** Phenotype of cell populations corresponding to sequential T-cell maturation stages in the human postnatal thymus analyzed for miRNA expression.

### SUPPLEMENTAL FIGURES AND LEGENDS

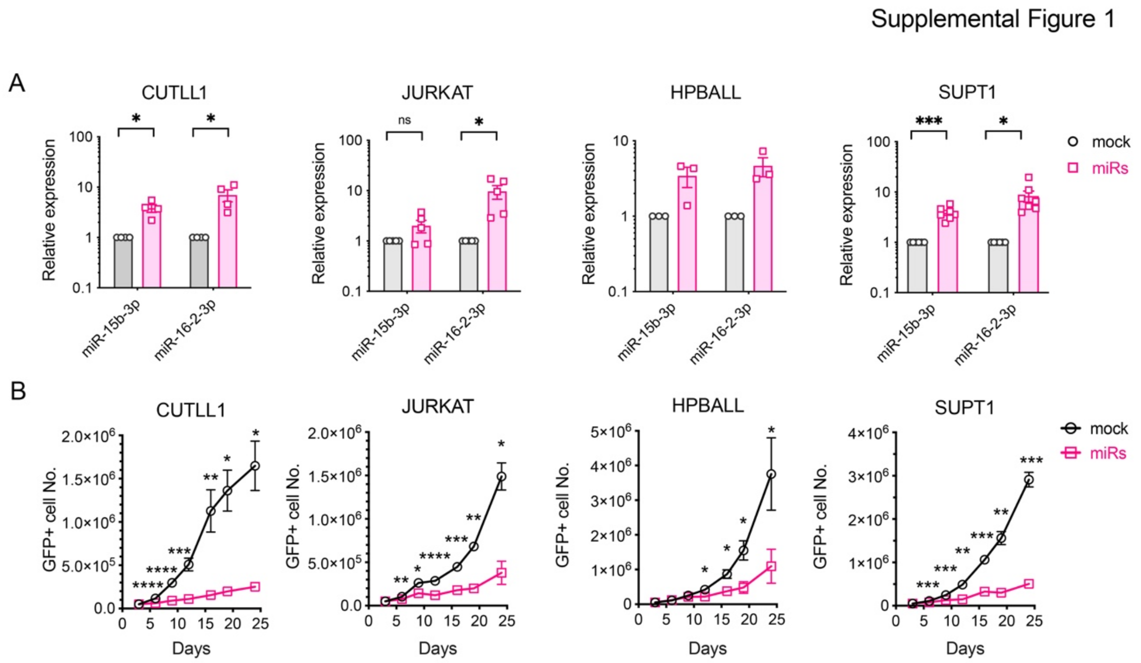


**Supplemental Figure 1. Ectopic expression of miR-15b/16-2 impairs *in vitro* expansion of human T-ALL cell lines.** (A) Relative expression of miR-15b-3p and miR-16-2-3p determined by RT-qPCR in CUTLL1, JURKAT, HPBALL and SUPT1 T-ALL cell lines transduced with either miR-15b/16-2 plus GFP (miRs), or GFP alone (mock) as control. RNAU44 was used as endogenous RT-qPCR control. Data are shown as mean relative expression (RQ) ± SEM of independent experiments run in triplicate normalized to mock-transduced cells (CUTLL1, n = 4; JURKAT, n = 5; HPBALL, n = 3; SUPT1, n = 7). Statistical differences were determined by one-sample t test. (B) Absolute numbers of T-ALL cells in (A) recovered upon *in vitro* culture at the indicated days post-transduction, normalized to 5 x $10^4$ initial cells. Data are shown as mean values ± SEM of independent experiments (n ≥ 3 for each T-ALL cell line). **P < .05; **P < .01; ***P < .001; ****P<.0001.*

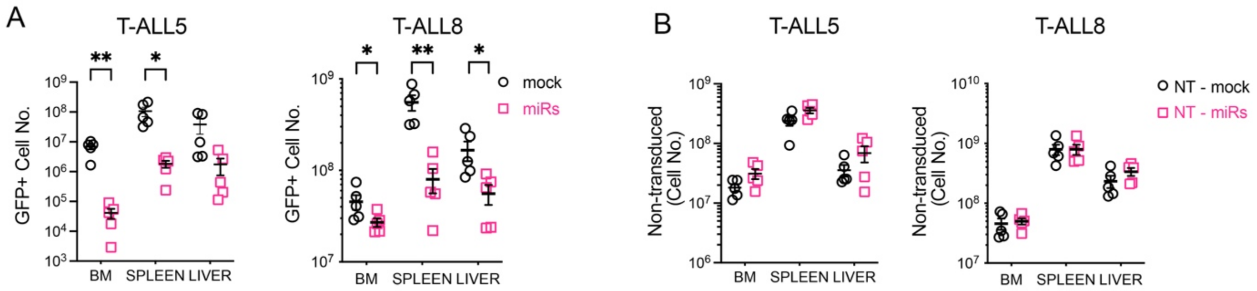


**Supplemental Figure 2. miR-15b/16-2 overexpression impairs *in vivo* progression of human T-ALL in patient-derived xenotransplantation assays.** (A) Absolute numbers of mock- and miR-15b/16-2-transduced primary T-ALL5 and T-ALL8 patient cells recovered from the indicated organs of xenotransplanted immunodeficient mice. Data are shown as mean values ± SEM normalized to $10^5$ mock- or miRs-transduced input injected cells. (B) Mean absolute numbers ± SEM of non-transduced (NT) T-ALL cells recovered from the indicated organs of immunodeficient mice in (A) normalized to $10^5$ NT input injected cells from mock- (NT-mock) or miR-15b/16-2-transduced (NT-miRs) T-ALL5 and T-ALL8 cell suspensions. Statistical differences were determined by Student's t tests. **P < .05; **P < .01.*

Supplemental Figure 3

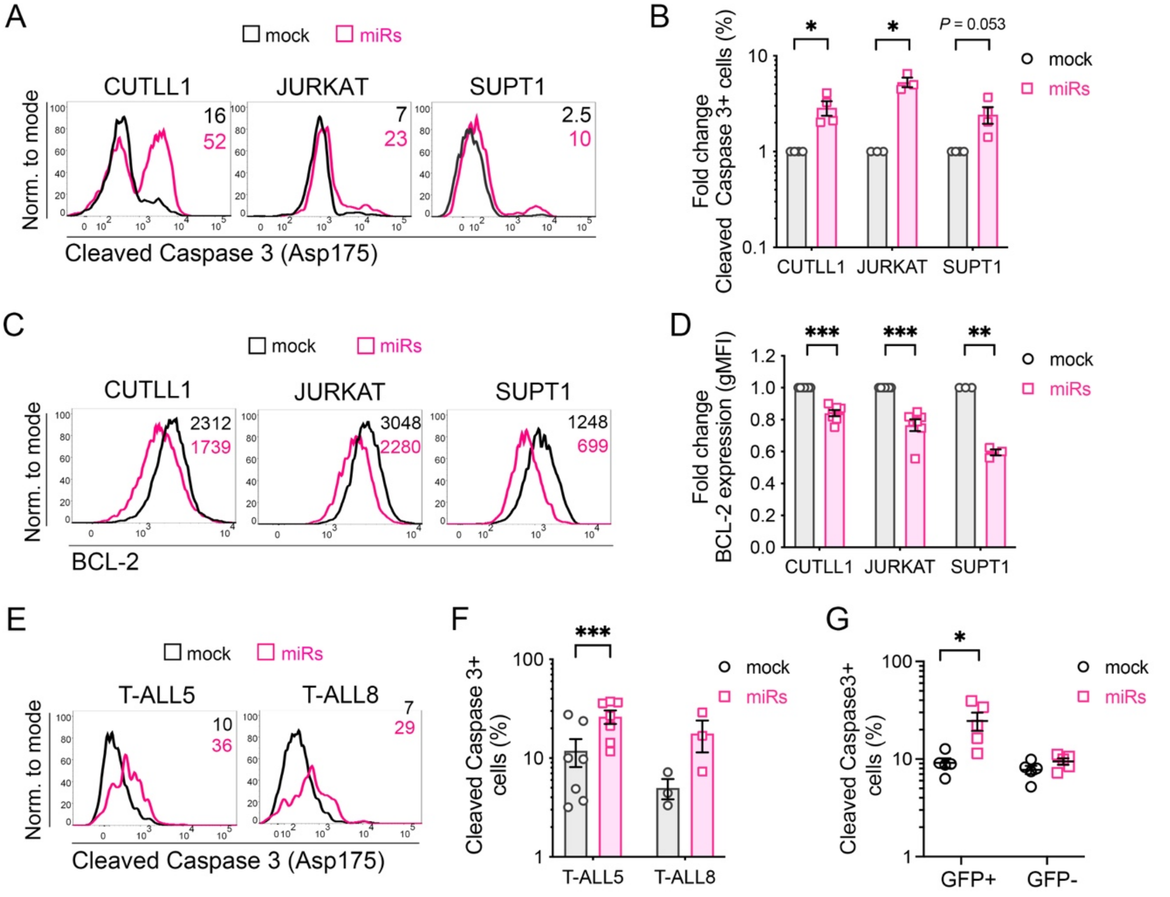


**Supplemental Figure 3. miR-15b/16-2 overexpression induces apoptosis in human T-ALL.** (A) Representative analysis of cell apoptosis determined by flow cytometry of cleaved Caspase 3 (Asp175) expression in human CUTLL1, JURKAT and SUPT1 T-ALL cell lines transduced with lentiviral vectors encoding either miR-15b/16-2 plus GFP (miRs) or GFP alone (mock) at day 3 post-transduction. Numbers are percentages of cleaved Caspase $3^+$ apoptotic cells. (B) Mean percentages ± SEM of apoptotic cells from different experiments (CUTLL1, n = 4; JURKAT, n = 3; SUPT1, n = 4) performed as in (A). Statistical differences were determined by one-sample t test. (C) Representative BCL-2 expression in miRs- and mock-transduced CUTLL1, JURKAT and SUPT1 human T-ALL cell lines determined by flow cytometry at day 3 post-transduction. Numbers represent the geometric mean of BCL-2 fluorescence intensity (gMFI). (D) Mean ± SEM of BCL-2 expression levels (gMFI) in miRs-transduced relative to mock-transduced cells in different experiments (CUTLL1, n = 7; JURKAT, n = 7; SUPT1, n = 3) performed as in (C). Statistical differences were determined by one-sample t test. (E) Representative flow cytometry analysis of cell apoptosis determined by flow cytometry of cleaved Caspase 3 (Asp175) expression in patient-derived T-ALL5 and T-ALL8 samples at day 3 post-transduction with lentiviral vectors encoding either miR-15b/16-2 plus GFP (miRs) or GFP alone (mock). Numbers are percentages of cleaved Caspase $3^+$ apoptotic cells. (F) Mean percentages ± SEM of apoptotic cells from different experiments (T-ALL5, n = 7; T-ALL8, n = 3) performed as in (E). (G) Mean percentages ± SEM of cleaved Caspase $3^+$ apoptotic cells among non-transduced ($GFP^-$), and mock- or miRs-transduced ($GFP^+$) T-ALL cells, recovered from the BM of immunodeficient mice at 12.5-weeks post-transplant with mock- or miRs-transduced T-ALL5 patient cells as shown in Figure 3 (n = 5 mice/group). Statistical differences in (F, G) were determined by Student's t tests. **P < .05; **P < .01; ***P < .001.*

Supplemental Figure 4

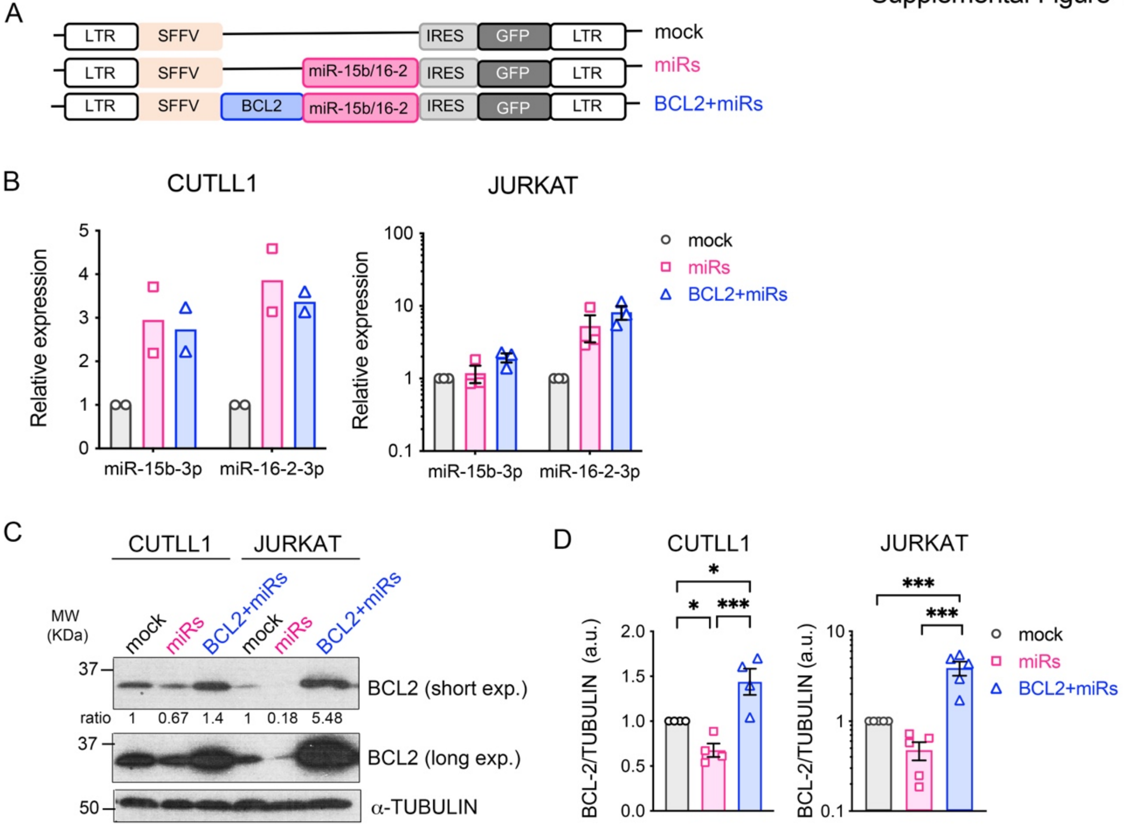


**Supplemental Figure 4. Generation of a BCL-2 rescue lentiviral vector.** (A) Schematic representation of a BCL-2 rescue vector encoding BCL-2, miR-15b/16-2 and GFP (BCL2 + miRs) used to test the ability of BCL-2 to restore miR-15b/16-2-mediated survival defects in T-ALL. Lentiviral vectors encoding either the miR-15b/16-2 cluster plus GFP as cell tracer (miRs) or GFP alone as control (mock) are also shown. (B) Relative expression of miR-15b-3p and miR-16-2-3p in CUTLL1 and JURKAT T-ALL cell lines transduced with the lentiviral vectors shown in (A), analyzed by RT-qPCR at day 10 post-transduction. RNAU44 was used as endogenous control. Data are mean of relative expression (RQ) ± SEM normalized to mock-transduced cells of independent experiments (CUTLL1, n = 2; JURKAT, n = 3) run in triplicate. (C) Representative immunoblot analysis of BCL-2 expression in CUTLL1 and JURKAT T-ALL cell lines transduced with the lentiviral vectors shown in (A). α-TUBULIN was used as protein loading control. Molecular weight (Mw) is indicated on the left. Two different exposure times (short and long) are shown. Numbers indicate the ratio between BCL-2 and α-TUBULIN expression, relative to mock-control transduced cells. (D) Quantification of BCL-2 expression levels relative to α-TUBULIN levels (a.u., arbitrary units) in different experiments (CUTLL1, n = 4; JURKAT, n = 5) performed as in (C). Data are shown as mean values ± SEM normalized to mock-transduced cells. Statistical differences were determined by one-way ANOVA with Tukey posttest analysis. *$^*P < .05$; $^{***}P < .001$.*

Supplemental Figure 5

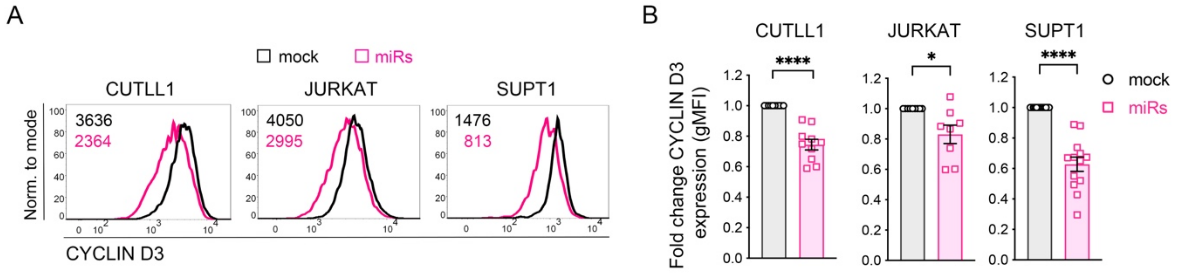


**Supplemental Figure 5. miR-15b/16-2 overexpression reduces CYCLIN D3 expression in human T-ALL.** (A) Representative CYCLIN D3 protein expression in T-ALL cell lines transduced with lentiviral vectors encoding either miR-15b/16-2 plus GFP (miRs) or GFP alone (mock), determined by flow cytometry at day 4 (CUTLL1, SUPT1), and day 7 (JURKAT) post-transduction. Numbers represent gMFI of CYCLIN D3 expression. (B) Quantification of CYCLIN D3 expression measured as gMFI in miRs-transduced relative to mock-transduced T-ALL cells. Data are mean values ± SEM of independent experiments performed as in (A) (CUTLL1, n = 10; JURKAT, n = 8; SUPT1, n =13). Statistical differences were determined by one sample t test. **P < .05; ****P < .0001.*
